# PD-L1^ATTAC^ mice reveal the potential of targeting PD-L1 expressing cells in cancer therapy

**DOI:** 10.1101/2022.07.22.501095

**Authors:** Elena Fueyo-Marcos, Coral Fustero-Torre, Gema Lopez, Marta Elena Antón, Fátima Al-Shahrour, Oscar Fernández-Capetillo, Matilde Murga

## Abstract

Antibodies targeting the PD-1 receptor and its ligand PD-L1 have shown impressive responses in some tumors of bad prognosis. We hypothesized that the selective elimination of PD-L1 expressing cells could similarly have antitumoral effects. To address this question, we developed an inducible suicidal knock-in mouse allele of *Pd-l1* (PD-L1^ATTAC^) which allows for the tracking and specific elimination of PD-L1-expressing cells in adult tissues. Elimination of PD-L1 expressing cells from the mouse peritoneum increased the septic response to lipopolysaccharide (LPS), due to an exacerbated inflammatory response to the endotoxin. In addition, mice depleted of PD-L1^+^ cells were resistant to colon cancer peritoneal allografts, which was associated with a loss of immunosuppresive B cells and macrophages, concomitant with an increase in activated cytotoxic CD8 T cells. Collectively, these results illustrate the usefulness of PD-L1^ATTAC^ mice and provide genetic support to the concept of targeting PD-L1 expressing cells in cancer therapy.

## Introduction

Activation of T-cells involves binding of the T cell receptor (TCR) on T cells to peptide-bound major histocompatibility complexes (MHC) on antigen presenting cells (APCs). This activating signal is modulated by membrane-bound co-stimulatory receptors that potentiate the response or by co-inhibitory receptors which limit self-damage (Smith-Garvin et al. 2009). The discovery of immune checkpoint mediated by PD-1 and CTLA-4 receptors, and that targeting these pathways potentiates the antitumoral capabilities of our immune system led to the Nobel Prize in Medicine in 2018 (Kroemer and Zitvogel 2021). Among all cancer immunotherapy strategies, antibodies targeting the PD-1/PD-L1 interaction have been the most intensively evaluated in clinical trials and are currently approved for a wide range of malignancies including melanoma, non-small cell lung cancer, Hodgkin’s lymphoma, head and neck squamous cell carcinoma or solid tumors presenting microsatellite instability (MSI) (Gong et al. 2018; Sun et al. 2018). Despite the undisputable success of this therapies, unfortunately only 20-40% of the patients respond to the therapy and even fewer show long-term responses (Garon et al. 2019; Hamid et al. 2019). In addition, resistance to therapy, intrinsic or acquired, is also a frequent clinical finding in patients treated with immune checkpoint inhibitors (ICIs) (Sharma et al. 2017). Consistently, current efforts are directed to identify strategies that increase the percentage of patients that respond to cancer immunotherapy, and the efficacy of these treatments.

We hypothesized that, since PD-L1 expressing APCs might display additional immune checkpoint mediators on their membranes, their elimination could also exert antitumoral potential. In fact, recent studies have reported that chimeric antigenic receptor T (CAR-T) cells targeting PD-L1 reduce the growth of solid tumor xenografts in mice (Xie et al. 2019; Qin et al. 2020). To provide genetic proof-of-principle support for the validity of this approach, we generated mice carrying an inducible suicidal reporter allele of PD-L1. Besides its usefulness to identify and isolate PD-L1 expressing cells (PD-L1^+^, hereafter) from adult mouse tissues, our work with these mice reveals that the selective elimination of PD-L1^+^ cells potentiates immune responses against different stimuli such as bacterial endotoxins or immunogenic cancer cells.

## Results and Discussion

### Generation of an inducible suicidal mouse model of PD-L1

To generate an inducible suicidal reporter allele of PD-L1, we used the previously developed ATTAC (apoptosis through targeted activation of caspase 8) strategy (Pajvani et al. 2005). In brief, we knocked in EGFP at the start codon of the mouse PD-L1 gene (*Cd274*), followed by FLAG-tagged catalytic domains of human caspase 8 fused to dimerizing serial FKBP domains which expression is driven by an IRES (**Fig. 1A**). This *PD-L1^ATTAC^* allele allows for the identification of PD-L1^+^ cells on the basis of EGFP expression, as well as their selective killing through apoptosis upon treatment with the FK102 analogue AP20187. After identifying successfully recombined mouse embryonic stem cell (mESC) clones by Southern Blotting (**Fig. 1B**) and before proceeding into generating mice, we first wanted to further confirm the correct integration of the allele by evaluating EGFP expression. To do so, we used the synergistic activation mediator (SAM) strategy, which enables CRISPR-dependent transcriptional activation of a selected gene (Konermann et al. 2015). Upon lentiviral transduction of *PD-L1^ATTAC^* heterozygous mESC (*PD-L1^AT/+^*) with an sgRNA targeting the *Cd274* promoter, EGFP expression was detected throughout the infected population (**Fig. 1C**). *PD-L1^AT/+^* mESC were then used to generate mice using standard procedures and crosses between *PD-L1^ATTAC^* heterozygous mice yielded *PD-L1^+/+^, PD-L1^AT/+^ and PD-L1^AT/AT^* animals at Mendelian rations. Mutant mice showed no apparent phenotype when compared to wild type (wt) littermates (**Fig. 1D**). However, and consistent with the fact that *PD-L1^AT/AT^* animals are knockouts for *Pdl1*, allografts of B16-F10 melanoma cells presented more immune infiltrates and grew less and when implanted in *PD-L1^AT/AT^* mice than in wt animals (**Fig. S1**).

**Figure 1.**
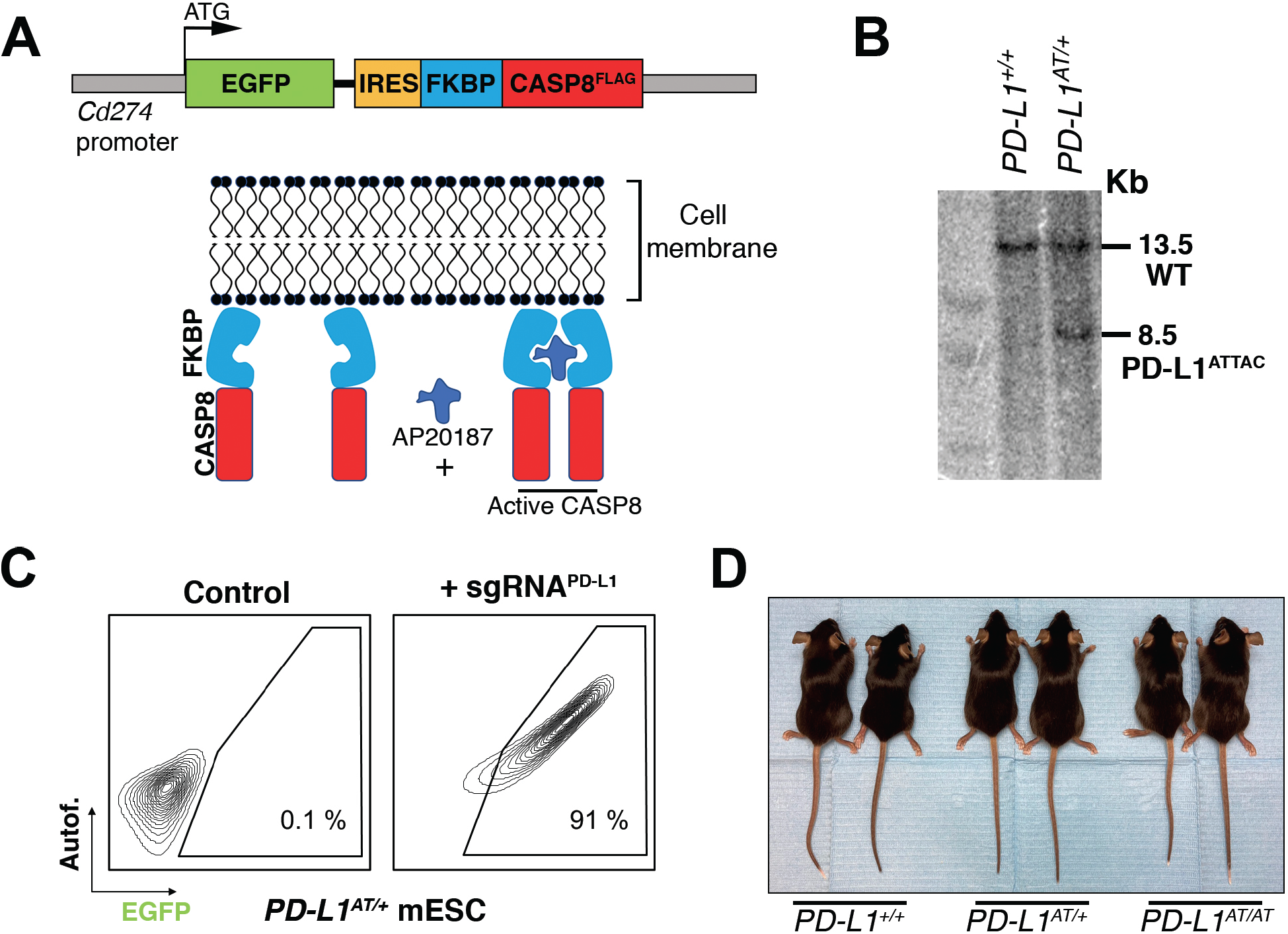
Generation of an inducible suicidal mouse model of PD-L1 (PD-L1^ATTAC^). (A) Scheme illustrating the construct used for the generation of PD-L1^ATTAC^ mice. The construct is under control of *Cd274* promoter and codes for EGFP as a reporter gene and a FLAG-tagged caspase 8 fused to FKBP domains, which homodimerize in the presence of AP20187 and induce apoptosis of PD-L1^+^ cells. (B) Southern blot illustrating the presence of mESC clones harboring the correct integration of the PD-L1^ATTAC^ allele that were subsequently used for the generation of mutant mice. The 13.2 Kb band corresponds to the endogenous WT *Cd274* gene, and the 8.5 Kb band to the knock-in PD-L1^ATTAC^ allele. (C) FACS analyses of PD-L1 expression as monitored by EGFP in PD-L1^ATTAC^ mESC cells harboring a dead Cas9 compatible with the SAM CRISPR activator system and transduced or not with a sgRNA against the *Cd274* promoter. Percentage of EGFP+ cells is shown. (D) Representative picture from pairs of 3-month-old animals of the indicated genotypes.

### In vitro validation of the PD-L1^ATTAC^ model

To evaluate the usefulness of the system *in vitro* we first generated mouse embryonic fibroblasts (MEF). Western Blotting (WB) revealed that a treatment with interferon gamma (IFNγ), a known inducer of PD-L1 expression (Kim et al. 2005; Lee et al. 2005), triggered expression of EGFP in *PD-L1^AT/+^* and *PD-L1^AT/AT^* but not in wt MEF (**Fig. 2A**). Conversely, PD-L1 expression was detectable in wt and *PD-L1^AT/+^* MEF but not in homozygous mutants, which is expected as the construct was inserted at the start codon and is thus a knockout allele (**Fig. 2A**). Equivalent findings were made by immunofluorescence (IF) (**Fig. 2B**) or flow cytometry (**Fig. 2C**). Moreover, analysis of flow cytometry data revealed a full correlation between EGFP and PD-L1 expression in IFNγ-treated MEF (**Fig. 2D**). In what regards to the cell killing induced by AP20187 and, to our surprise, we could only detect a significant impact in cell death if *PD-L1^AT/+^* or *PD-L1^AT/AT^* MEF were previously treated with IFNγ to trigger PD-L1 expression and also grown in low serum media (0.1% FBS) (**Fig. 2E**). In contrast, AP20187 failed to significantly induce cell death if MEF were grown in media containing 10% FBS (**Fig. 2F**). We hypothesized that this could be due to the ATTAC system being particularly efficient in killing non-growing cells as these cells might have a lower threshold for triggering apoptosis. In support of this view, we want to note that despite the usefulness of this strategy it has only been previously used to target non-dividing cells such as adipocytes, pancreatic islet beta cells or senescent cells (Pajvani et al. 2005; Wang et al. 2008; Baker et al. 2011).

**Figure 2.**
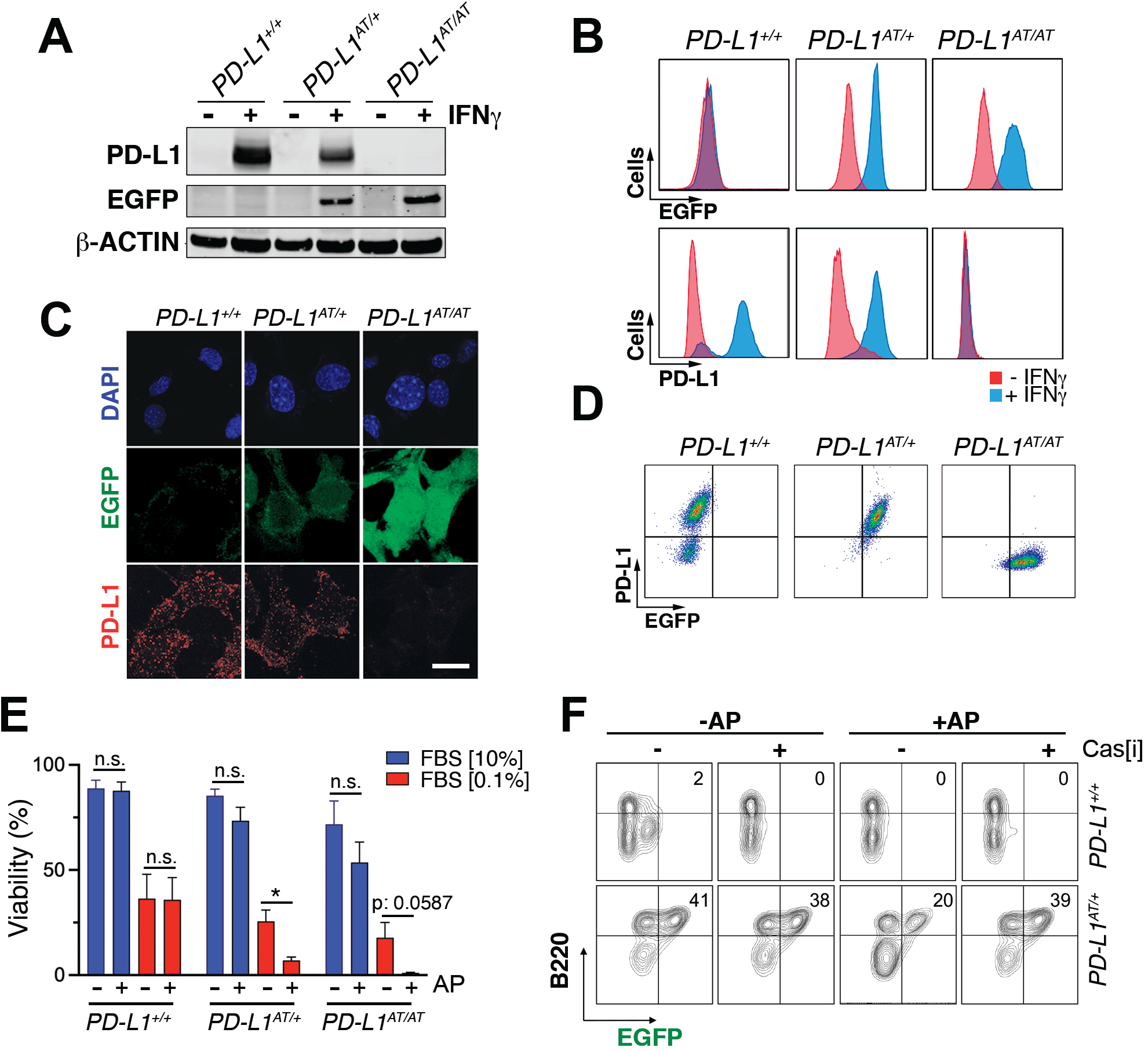
*In vitro* validation of the PD-L1^ATTAC^ mouse model. (A) Western blot illustrating PD-L1 and EGFP expression in *PD-L1*^+/+^, *PD-L1*^AT/+^ and *PD-L1*^AT/AT^ MEFs exposed or not to IFNγ (100 ng/ml) for 48 hours. (B,C) Immunofluorescence (B) and flow cytometry (C) analyses of EGFP and PD-L1 expression in *PD-L1*^+/+^, *PD-L1*^AT/+^ and *PD-L1*^AT/AT^ MEFs exposed to IFNγ (100 ng/ml) for 48 hours. Scale bar (white) indicates 5 μm. (D) Two-dimensional dot plot from the flow cytometry data shown in (C) illustrating the correlation between EGFP and PD-L1 expression per cell. (E) Percentage of live cells by FACS in *PD-L1*^+/+^, *PD-L1*^AT/+^ and *PD-L1*^AT/AT^ MEFs cultured in normal or low-serum media (0.1% FBS) containing IFNγ (10 ng/ml) and treated or not with AP20187 (100 nM). Cells were cultured in normal or low-serum media for 24 hours. The day after, cells were exposed or not to AP20187 for 72 hours. (F) FACS analyses of B220 and EGFP expression of splenocytes from *PD-L1*^+/+^ and *PD-L1*^AT/+^ mice cultured in IFNγ (10 ng/ml), LPS (10 ng/ml) and M-CSF (10 ng/ml) for 24 hours before exposition to AP20187 (100 nM) and caspase inhibitor I (20 μM) for 24 hours. Percentage of B220+ EGFP+ cells is shown.

Next, and given that immune cells are the main source of PD-L1 *in vivo*, we isolated splenocytes from adult mice, stimulated B cells with LPS and treated the cultures with IFNγ to induce PD-L1 expression. These experiments revealed a clear population of EGFP positive B cells (identified on the basis of B220 expression), which was selectively killed upon treatment with AP20187 (**Fig. 2E**). Moreover, this cell killing could be prevented with a pan-caspase inhibitor, confirming that cell death was due to apoptosis (**Fig. 2E**). Collectively, these data indicate that the *PD-L1^ATTAC^* allele is efficient for the tracing of PD-L1^+^ cells, as well as for enabling their clearance in several cell types such as LPS-stimulated B cells or MEF.

### In vivo validation of the PD-L1^ATTAC^ model

To evaluate the usefulness of the *PD-L1^ATTAC^* allele as a reporter of PD-L1 expression in vivo, we first stained tissues of adult mice with an anti-EGFP antibody. As expected, EGFP expression was highest in organs from the immune system such as the spleen, lymph nodes, bone marrow or the thymus. In addition, scattered expression could also be detected in other organs such as the intestine, lungs or liver, while no expression was seen in the kidneys, pancreas, or hearts (**Fig. 3A and Fig S2A**). Similar observations were also made by WB (**Fig. 3B**). Importantly, dual staining in lungs from wt and *PD-L1^AT/+^* animals identified that cells with cytoplasmic EGFP expression also presented PD-L1 on their membranes (**Fig. 3C**), further supporting the reporter nature of the introduced mutation. Furthermore, while EGFP expressing cells were readily seen in homozygous *PD-L1^AT/AT^* lungs, these cells lacked PD-L1 expression, consistent with the null nature of the allele (**Fig. 3C**). Immunofluorescence experiments further identified cells expressing both PD-L1 and EGFP in lungs from *PD-L1^AT/+^* mice (**Fig. 3D**).

**Figure 3.**
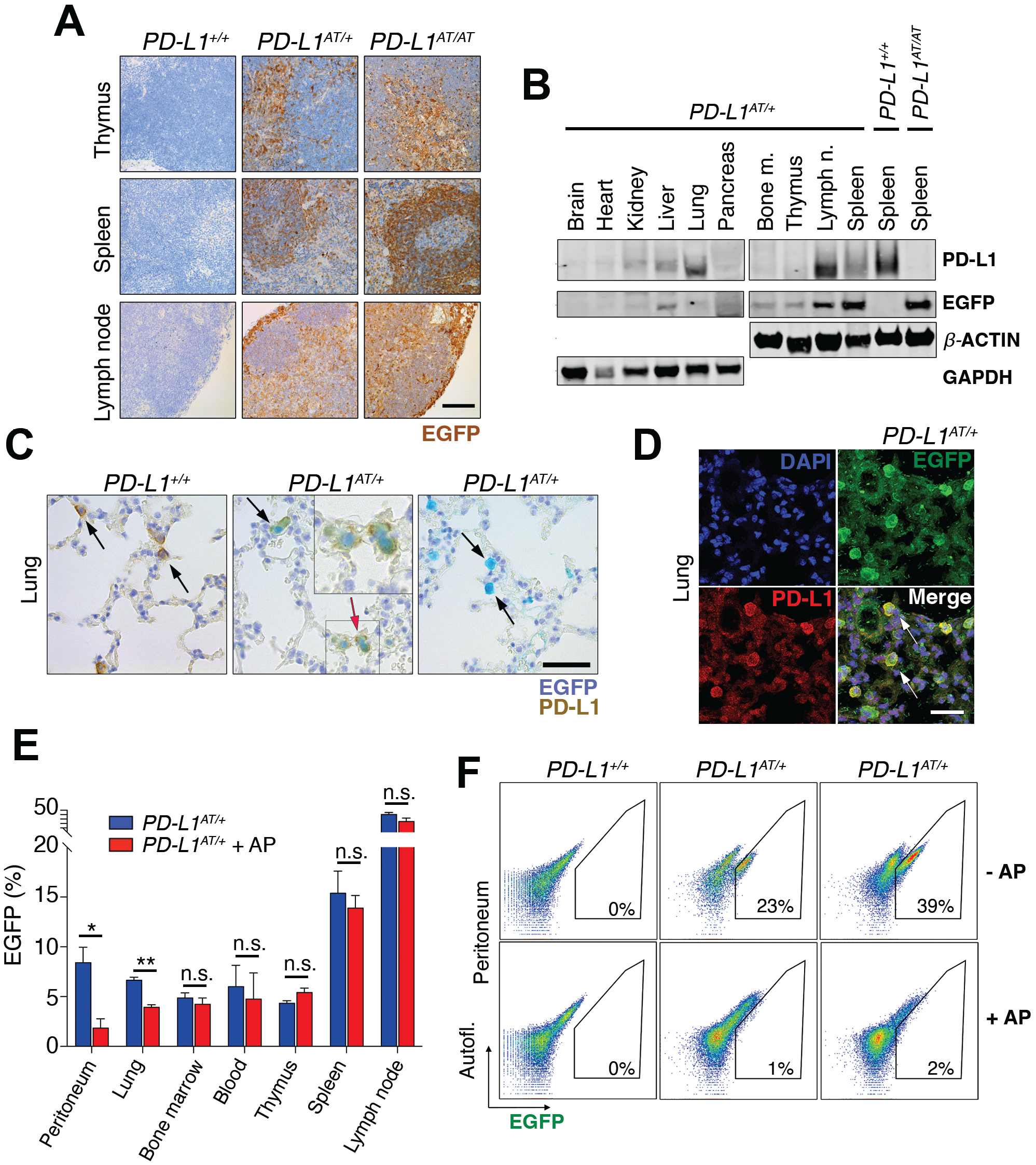
*In vivo* validation of the PD-L1^ATTAC^ mouse model. (A) EGFP immunohistochemistry (IHC) from the thymus, spleen and lymph nodes of *PD-L1*^+/+^, *PD-L1*^AT/+^ and *PD-L1*^AT/AT^ mice. Scale bar (black) indicates 100 μm. (B) Western blot illustrating PD-L1 and EGFP expression in different organs of *PD-L1*^AT/+^ mice and the spleen of wt and *PD-L1*^AT/AT^ mice. Actin and GAPDH were used as a loading control. (C) Representative images from a dual PD-L1 and EGFP IHC in lungs from *PD-L1*^+/+^, *PD-L1*^AT/+^ and *PD-L1*^AT/AT^ mice. Arrows indicate examples EGFP expressing cells. The red arrow in the *PD-L1*^AT/+^ panel indicates an inset that is magnified in the right-hand corner to illustrate the appearance of cells expressing both EGFP and PD-L1. Scale bar (black) indicates 50 μm. (D) Representative image from a dual EGFP and PD-L1 IF in the lung of *PD-L1*^AT/+^. Scale bar (white) indicates 30 μM. (E) Percentage of EGFP+ cells as revealed by FACS in different organs from control and AP20187-treated (AP) *PD-L1*^AT/+^ mice. AP20187 was administered via I.P. at 2.5 mg/kg for 3 days. The p value was calculated with unpaired t-test. n.s.: non-significant, *p < 0.05. *p < 0.01 (F) FACS analysis of PD-L1 expression as monitored by EGFP in peritoneal cells from *PD-L1*^+/+^, *PD-L1*^AT/+^ and *PD-L1*^AT/AT^ mice treated or not with AP20187 (2.5 mg/kg) for 3 days. Percentage of EGFP+ cells is shown.

In what regards to the inducible-suicidal properties, we first evaluated the effects of an intraperitoneal (i.p.) administration of AP20187 in *PD-L1^ATTAC^* mice. The treatment was particularly efficacious in killing EGFP+ cells in the peritoneum, although we also saw a significant effect in the lungs (**Fig. 3E**). In contrast, this approach had no significant impact in reducing the percentage of EGFP expressing cells in the blood, thymus, spleen or lymph nodes (**Fig. 3E**). FACS analyses confirmed a very efficient depletion of EGFP expressing cells from the peritoneum of *PD-L1^AT/+^* and *PD-L1^AT/AT^* mice after treatment with AP20187 (**Fig. 3F**). We also evaluated if an intravenous (i.v.) administration of the drug could have more widespread effects. However, while i.v. delivery of AP20187 led to a significant depletion of EGFP+ cells in the blood and bone marrow, we failed to see significant effects on other tissues (**Fig. S2B**). On the basis of these results, we decided to focus in the adult peritoneum as a model where to study the impact of selectively targeting PD-L1^+^ cells.

### Depletion of PD-L1 expressing cells sensitizes mice to LPS

Intraperitoneal injection of the bacterial lipopolysaccharide (LPS) is a widely used experimental model of a lethal septic shock associated to a cytokine storm (Redl et al. 1998). Strikingly, while i.p. injections of AP20187 for three days did not affect LPS-mortality in wild type mice, it led to a significant reduction of the survival of *PD-L1^AT/+^* animals (**Fig. 4A,B**). This effect was even more pronounced in *PD-L1^AT/AT^* mice with all animals dying by 18hrs after LPS injection (with no wt animals being dead at this timepoint) (**Fig. 4C**). Consistent with survival data, treatment of *PD-L1^AT/+^* and *PD-L1^AT/AT^* mice with AP20187 triggered a higher accumulation of the inflammatory cytokine IL-6 in the plasma of LPS-injected mice, confirming the increased severity of the septic shock (**Fig. S3A-C**). Moreover, immunohistochemistry (IHC) analyses revealed a clear accumulation of immune infiltrates in the lungs or livers from AP20187-treated *PD-L1^AT/+^* and *PD-L1^AT/AT^* mice exposed to LPS (**Fig. 4D** and **Fig. S3D**). Together, these data illustrate that the selective elimination of PD-L1^+^ cells increases the severity of immune responses in the mouse peritoneum.

**Figure 4.**
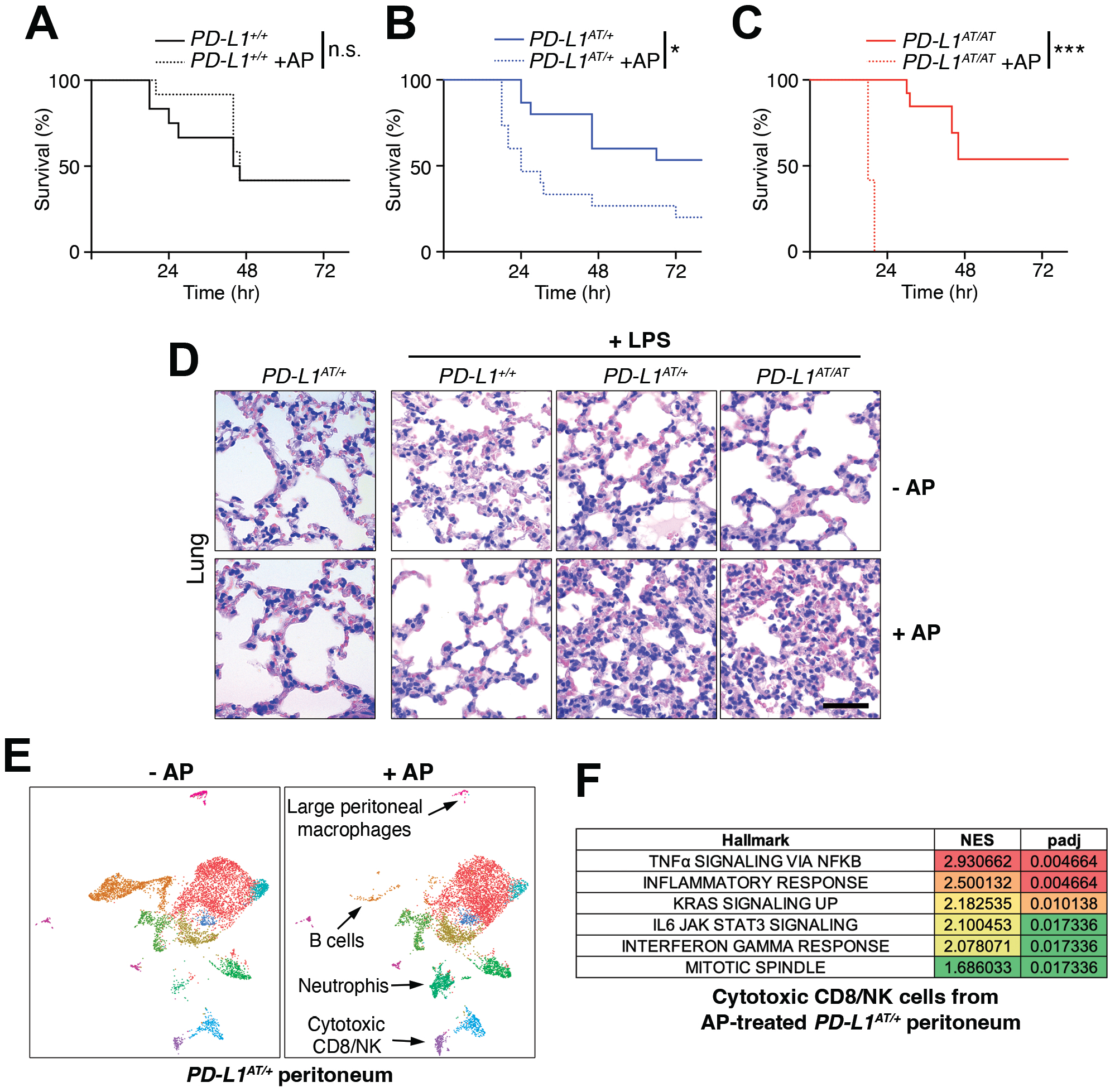
Effects of depleting PD-L1-expressing cells in a model of LPS-induced septicaemia. (A-C) Kaplan-Meier survival curves of *PD-L1*^+/+^, *PD-L1*^AT/+^ and *PD-L1*^AT/AT^ mice after LPS injection. Mice were treated via i.p. with AP20187 (2.5 mg/kg) for 3 days and subsequently injected i.p. with 10 mg/kg LPS. The p value was calculated with the Mantel-Cox log rank test. * p < 0.05 ***p < 0.001. (D) Hematoxylin/eosin IHC in the lungs from the experiment defined in (A-C). Note the further accumulation of infiltrates in the lungs of AP20187-treated *PD-L1*^AT/+^ and *PD-L1*^AT/AT^ mice after LPS injection. Scale bar (black) indicates 75 μm. (E) Single-cell sequencing analysis of the impact of AP20187 treatment (2.5 mg/kg, 3 days) on the repertoire of peritoneal cells from *PD-L1*^AT/+^ mice. Panels show UMAP plots from these analyses are shown and the cell types showing alterations are indicated by arrows. (F) GSEA analysis showing the hallmarks that were most significantly upregulated in cluster 9 (cytotoxic CD8/NK cells) after AP20187 treatment from the experiment defined in (E).

To determine which changes in cell types were responsible for the observed effects, we conducted single cell RNA sequencing analyses of intraperitoneal cells from untreated or AP20187-treated *PD-L1^AT/+^* mice. These analyses revealed a drug-induced depletion of B cells and macrophages, concomitant to an accumulation of neutrophils and cytotoxic CD8/NK cells (**Fig. 4E** and **Fig. S3E**). Moreover, Gene Set Enrichment Analyses (GSEA) indicate that CD8 cells from AP20187-treated *PD-L1^AT/+^* mice were hyperactivated, as revealed by the significant activation of several pathways such as those related to TNFα, IFNγ or IL-6 signaling, as well as a general activation of the inflammatory response (**Fig. 4F** and **Fig. S3F,G**). Hence, depletion of PD-L1^+^ cells alters the intraperitoneal immune repertoire which includes the accumulation of activated cytotoxic CD8 cells.

### Depletion of PD-L1+ cells increases survival to peritoneal tumor allografts

Finally, and given that cytotoxic T cells are thought to be the main effectors in the context of anti-PD-L1 immunotherapies (Tumeh et al. 2014), we tested the impact of depleting PD-L1^+^ cells in of cancer. To this end, we used a model of intraperitoneal metastasis by the highly immunogenic colon adenocarcinoma cell line MC-38, which can be used for allografts in immunocompetent mice and is sensitive to anti-PD-L1 therapies (Juneja et al. 2017). Furthermore, dissemination of cancer cells into the peritoneum is frequent in digestive and gynecological cancers and is associated with poor prognosis (Mikula-Pietrasik et al. 2018), highlighting the relevance of the chosen model.

To conduct these experiments, we first generated a MC-38 clone harboring constitutive expression of firefly luciferase which enables monitoring tumor progression by intravital imaging (MC-38^luc^). Intraperitoneal injection of MC-38^luc^ cells led to a lethal disease associated to the dissemination and growth of cancer cells in the peritoneal cavity (**Fig. 5A**). Treatment with AP20187 did not affect the progression of the disease in wt mice (**Fig. S4A**). Strikingly, while all *PD-L1^AT/+^* mice were dead within the first month after being intraperitoneally injected with MC-38^luc^ cells, a prior treatment with AP20187 prior to the injection of MC-38 cells led to survival of half of the animals (**Fig. 5B**). Consistently, intravital imaging showed a clear reduction of MC-38 cells in the peritoneum of AP20187-treated *PD-L1^AT/+^* mice (**Fig. 5C**). Furthermore, and similar to our previous observations in the LPS-sepsis model, intraperitoneal tumors from AP20187-treated presented a significant accumulation of infiltrating CD8 lymphocytes, highlighting the increased immune response to the tumor (**Fig. 5D**). Besides this prevention model, AP20187 treatment also increased the survival of *PD-L1^AT/+^* mice if the drug was administered 4 days after the injection of MC-38 cells (treatment model) (**Fig. S4B-D**). Together with the data from the LPS-induced septic shock, these experiments indicate that the selective targeting of PD-L1^+^ cells intensifies immune responses in the peritoneum, which in the context of cancer is protective and favours the clearance of tumor cells.

**Figure 5.** Depleting PD-L1^+^ cells prolongs survival in an immunocompetent model of peritoneal cancer metastasis. (A) Schematic overview of the prevention experimental workflow. 5·10^5^ MC-38^luc^ cells were intraperitoneally injected into mice that were previously injected i.p. for 3 consecutive days with AP20187 (2.5 mg/kg). (B) Kaplan-Meier survival curve of control and AP20187-pretreated *PD-L1*^AT/+^ mice after i.p. inoculation of MC-38^luc^ allografts. The p value was calculated with the Mantel-Cox log rank test. ***p < 0.001. (C) Representative IVIS image of mice from the experiment defined in (B) at day 4 post-tumour injection. (D) IHC of CD8 in intraperitoneal MC-38^luc^ allografts isolated from control and AP20187-treated *PD-L1*^AT/+^ mice. Insets in each panel are magnified to illustrate the presence of tumor-infiltrating CD8^+^ cells. Scale bar (black) indicates 30 μm.

In summary, we here present PD-L1^ATTAC^ as a reporter and inducible suicidal allele of mouse PD-L1. The reporter EGFP enables the identification and isolation of PD-L1+ cells from adult tissues, which we believe is useful as antibodies detecting mouse PD-L1 often give signal in PD-L1 deficient samples. As for the inducible-suicidal strategy, our experiments indicate that this effect is significantly influenced by the administration route for AP20187 and growth rates of the cells, which we wonder to what extent could also have influenced previous studies using the ATTAC strategy. Despite these limitations, i.p. administration of AP20187 yields a very efficient depletion of PD-L1^+^ from the peritoneum, enabling functional studies to investigate the impact of depleting PD-L1^+^ cells in vivo. Of note, no significant toxicities were observed in mice that were i.p. treated for up to 3 months with 3 weekly doses of AP20187. Our results confirm that the selective elimination boosts immune responses in the peritoneum, which prolongs survival against a model of peritoneal cancer metastasis. Hence, our work supports the usefulness of targeting PD-L1 expressing cells in cancer therapy, and provides the immunotherapy research community with a useful genetic tool for investigations on the PD-1/PD-L1 checkpoint in mice.

## Materials and Methods

### Mouse models

The PD-L1^ATTAC^ targeting vector was generated by recombineering (Genebridges, Germany). Recombinant ES cells were screened by Southern Blot through standard procedures, and subsequently used for the generation of chimaeras by aggregation. Animals were genotyped by PCR using the following primers (UP: 5’ - TTGCTTCAGTTACAGCTGGCTCG - 3’; Down_WT: 5’ - CGTAGCAAGTGACAGCAGGCTG - 3’; Down_MUT: 5’ – GCCGTTTACGTCGCCGTCCAG – 3’). Mice were kept under standard conditions at specific-pathogen free facility of the Spanish National Cancer Centre in a mixed C57BL/6-129/Sv background. 9–12-week-old mice were used for all experiments. All mouse work was performed in accordance with the Guidelines for Humane Endpoints for Animals Used in Biomedical Research, and under the supervision of the Ethics Committee for Animal Research of the “Instituto de Salud Carlos III”.

### AP20187 treatments

AP20187 (MedChemExpress, HY-13992) was resuspended in 75% EtOH at 62.5 mg/ml and stored at −20°C. For in vitro experiments, cells were treated with 100 nM AP20187 for 48-72h. In vivo, the compound was dissolved in 2% Tween-20 (Sigma, P7949) and 10% PEG-300 (Aldrich #202371) and mice were i.p. injected with 2.5 mg/kg AP20187.

### LPS-induced septicemia

Mice were injected i.p. with AP20817 (2.5 mg/kg) for 3 consecutive days before an i.p. injection of LPS (10 mg/kg, Sigma, #L2630) resuspended in PBS. Blood, plasma and tissues were isolated for further IHC and ELISA analyses. Mice were monitored for overall health and for a week after LPS injection.

### MC-38^luc^ allografts

Mice were injected i.p. with 5·10^5^ MC-38^luc^ cells. Tumor growth was monitored twice a week by intravital imaging using an IVIS Optica Imaging system (Perkin Elmer) after anesthetizing mice with Isoflurane. For IVIS analyses, animals were previously injected i.p. with 150 mg/kg D-Luciferin (Xenolight, #122799).

#### Statistics

All statistical analyses were performed using Prism software (GraphPad Software) and statistical significance was determined where the p-value was <0.05 (*), <0.01 (**), <0.001 (***) and <0.0001 (****). Survival data was evaluated using Kaplan-Meier analyses.

Extended Methods for this manuscript can be found in the Supplementary Materials.

## Acknowledgments

O.F-C. is supported by grants from the Spanish Ministry of Science, Innovation and Universities (RTI2018-102204-B-I00 and SAF2017-90900-REDT, co-financed with European FEDER funds) and the Spanish Association Against Cancer (AECC; PROYE20101FERN) to O.F-C. and by a Ph.D. fellowship from María Oliva-Amigos/as del CNIO to E.F-M. CNIO Bioinformatics Unit (BU) is a member of the Spanish National Bioinformatics Institute (INB), ISCIII-Bioinformatics platform and is supported by grant PT17/0009/0011, of the Acción Estratégica en Salud 2013-2016 of the Programa Estatal de Investigación Orientada a los Retos de la Sociedad, funded by the ISCIII and European Regional Development Fund (ERDF-EU) and project RETOS RTI2018-097596-B-I00 funded by AEI-MCIU and cofounded by the ERDF-EU. The authors declare no competing financial interests.

## Author Contributions

E.F. contributed to most of the experiments; C.F. and F.A.-S. helped with bioinformatic analyses of scRNAseq data; M.A. provided technical help; O.F. helped to supervise the work and to write the manuscript; M.M. coordinated the study, contributed to experiments and helped with manuscript writing.

