## Supplementary Figures and Methods for "PD-L1^ATTAC^ mice reveal the potential of targeting PD-L1 expressing cells in cancer therapy"

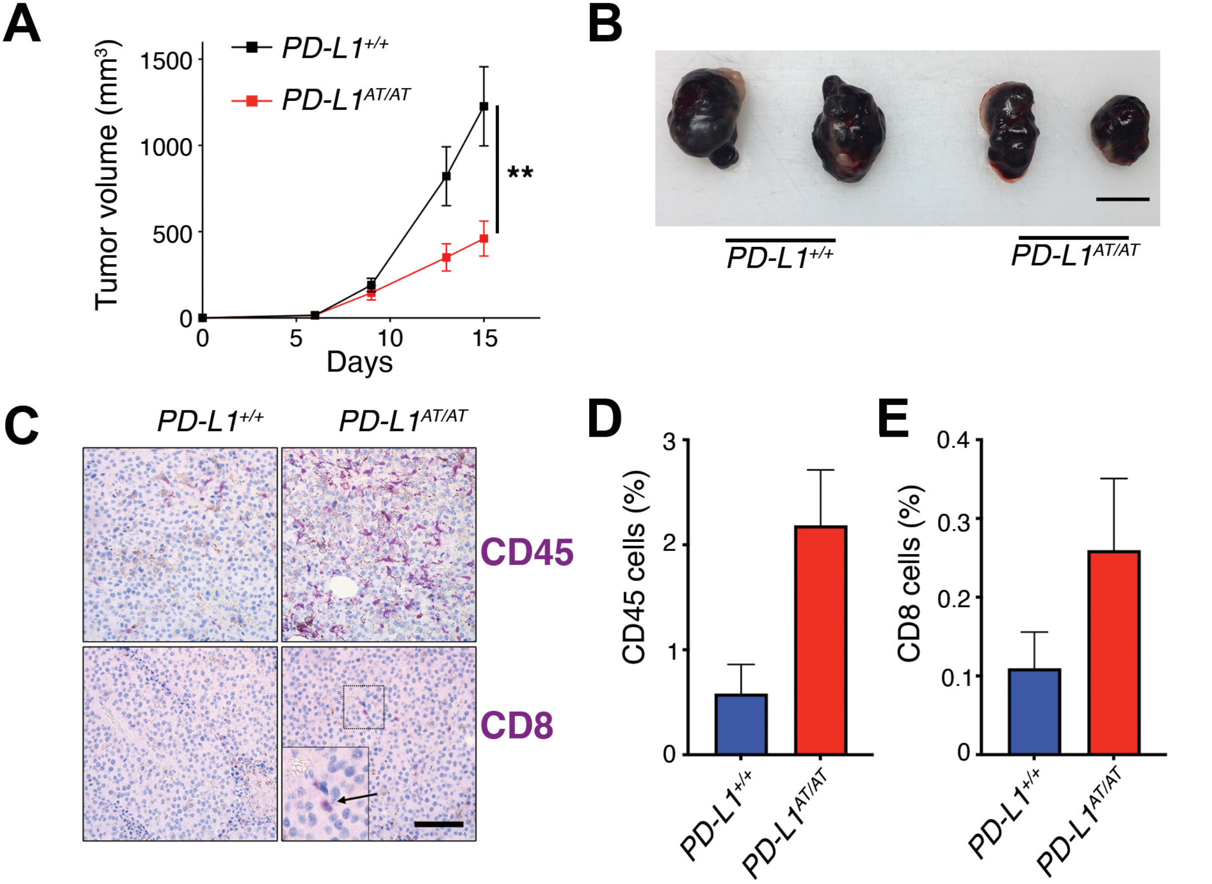
A
PD-L1+/+
PD-L1AT/AT
Tumor volume (mm3)
1500
1000
500
0
0
5
10
15
Days
**
B
PD-L1+/+
PD-L1AT/AT
C
PD-L1+/+
PD-L1AT/AT
CD45
CD8
D
CD45 cells (%)
3
2
1
0
PD-L1+/+
PD-L1AT/AT
E
CD8 cells (%)
0.4
0.3
0.2
0.1
0
PD-L1+/+
PD-L1AT/AT

**Figure S1. Resistance to B16-F10 melanoma allografts in *PD-L1*$^{AT/AT}$ mice.** (A) Growth of B16-F10 allografts subcutaneously implanted into the left flank of *PD-L1*$^{+/+}$ and *PD-L1*$^{AT/AT}$ mice. The p value was calculated with mixed-effects analysis. **p < 0.01 (B) Representative picture of the melanoma allografts from these analyses isolated at day 15. Scale bar (black) indicates 1 cm. (C) Immunohistochemistry of CD45 and CD8 in B16-F10 allografts isolated at day 15 from of *PD-L1*$^{+/+}$ and *PD-L1*$^{AT/AT}$ mice. An inset is magnified to illustrate the presence of tumor-infiltrating CD8+ cytotoxic T cells in the allografts grown in mutant mice. Scale bar (black) indicates 100 µm. (D,E) Quantification of CD45+ (D) and CD8+ (E) cells from the analyses shown in (C).

Fueyo-Marcos *et al.* **Fig S2**

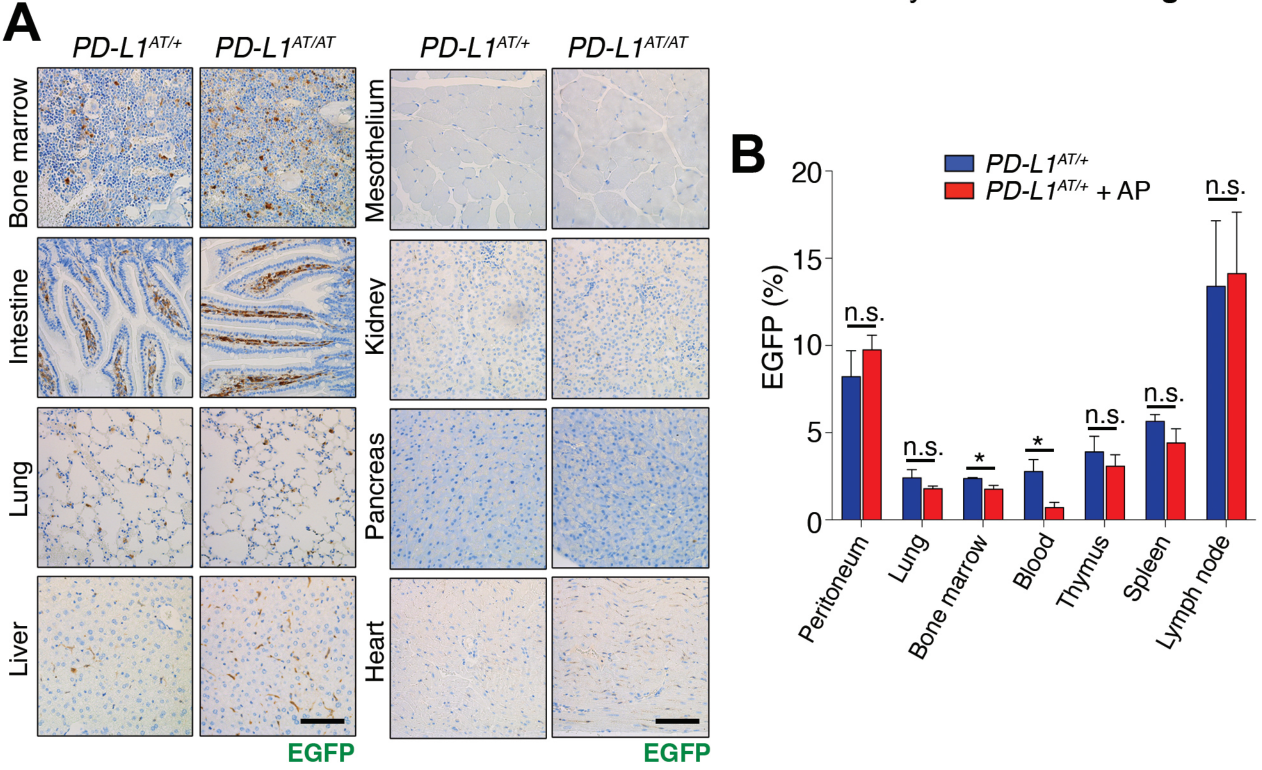

**Figure S2. Characterization of the PD-L1$^{ATTAC}$ mouse model**. (A) EGFP immunohistochemistry (IHC) from the bone marrow, intestine, lung, liver, mesothelium, kidney, pancreas and heart of *PD-L1*$^{AT/+}$ and *PD-L1*$^{AT/AT}$ mice. Scale bar (black) indicates 100 $\mu$m. (B) Percentage of EGFP+ cells as revealed by FACS in the indicated organs from control and AP-treated *PD-L1*$^{AT/+}$ mice. AP20187 (2.5mg/kg) was administered via i.v. for three consecutive days. The p value was calculated with unpaired t-test. n.s.: non-significant, *$p < 0.05$.

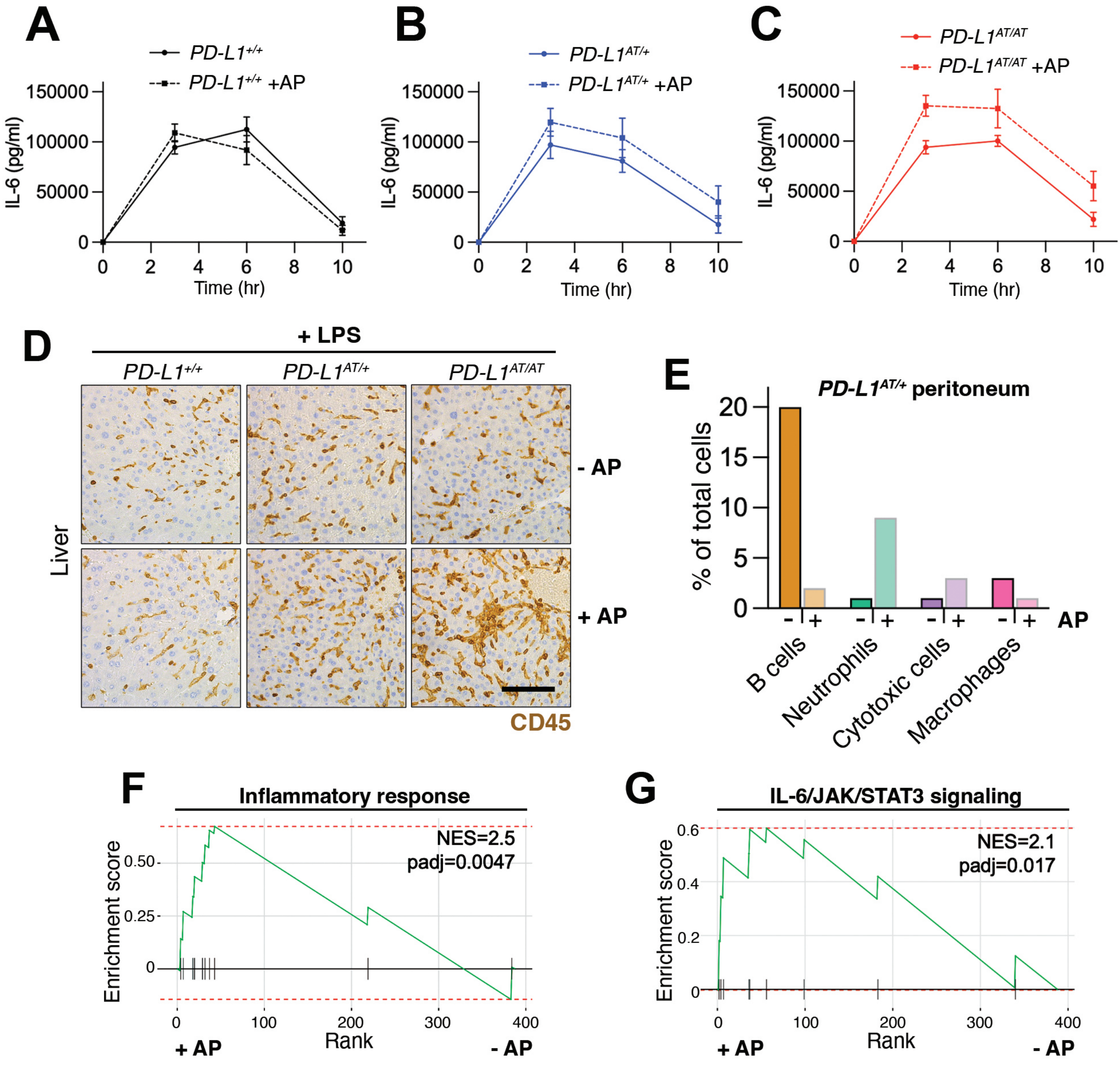
A
PD-L1+/+
PD-L1+/+ +AP
IL-6 (pg/ml)
150000
100000
50000
0
0 2 4 6 8 10
Time (hr)
B
PD-L1AT/+
PD-L1AT/+ +AP
IL-6 (pg/ml)
Time (hr)
C
PD-L1AT/AT
PD-L1AT/AT +AP
IL-6 (pg/ml)
Time (hr)
D
+ LPS
PD-L1+/+
PD-L1AT/+
PD-L1AT/AT
Liver
- AP
+ AP
CD45
E
PD-L1AT/+ peritoneum
% of total cells
20
15
10
5
0
- + - + - + - + AP
B cells
Neutrophils
Cytotoxic cells
Macrophages
F
Inflammatory response
NES=2.5
padj=0.0047
Enrichment score
0.50
0.25
0
0 100 200 300 400
Rank
+ AP
- AP
G
IL-6/JAK/STAT3 signaling
NES=2.1
padj=0.017
Enrichment score
0.6
0.4
0.2
0
0 100 200 300 400
Rank
+ AP
- AP

**Figure S3. Depletion of PD-L1$^{+}$ cells sensitizes mice to LPS.** (A-C) Kinetics of IL-6 accumulation and clearance in plasma isolated from *PD-L1*$^{+/+}$*, PD-L1*$^{AT/+}$ and *PD-L1*$^{AT/AT}$ mice after LPS injection as determined by ELISA. (D) IHC of CD45 in the livers of *PD-L1*$^{+/+}$*, PD-L1*$^{AT/+}$ and *PD-L1*$^{AT/AT}$ mice after LPS injection. Mice were treated via i.p. with AP (2.5 mg/kg) for 3 days and i.p. with 10 mg/kg LPS on the following day. Scale bar (black) indicates 100 μm. (E) Quantification from the single-cell sequencing analysis shown in **Fig. 4E**, indicating the changes in the cell repertoire in the peritoneum from *PD-L1*$^{AT/+}$ mice upon AP treatment. (F,G) Preranked GSEA on the genes from the hallmarks “Inflammatory response” (F) and “IL-6/JAK/STAT3 signaling” (G) obtained from scRNAseq data comparing the transcriptomes of cytotoxic cells from *PD-L1*$^{AT/+}$ mice upon AP treatment.

Fueyo-Marcos *et al.* **Fig S4**

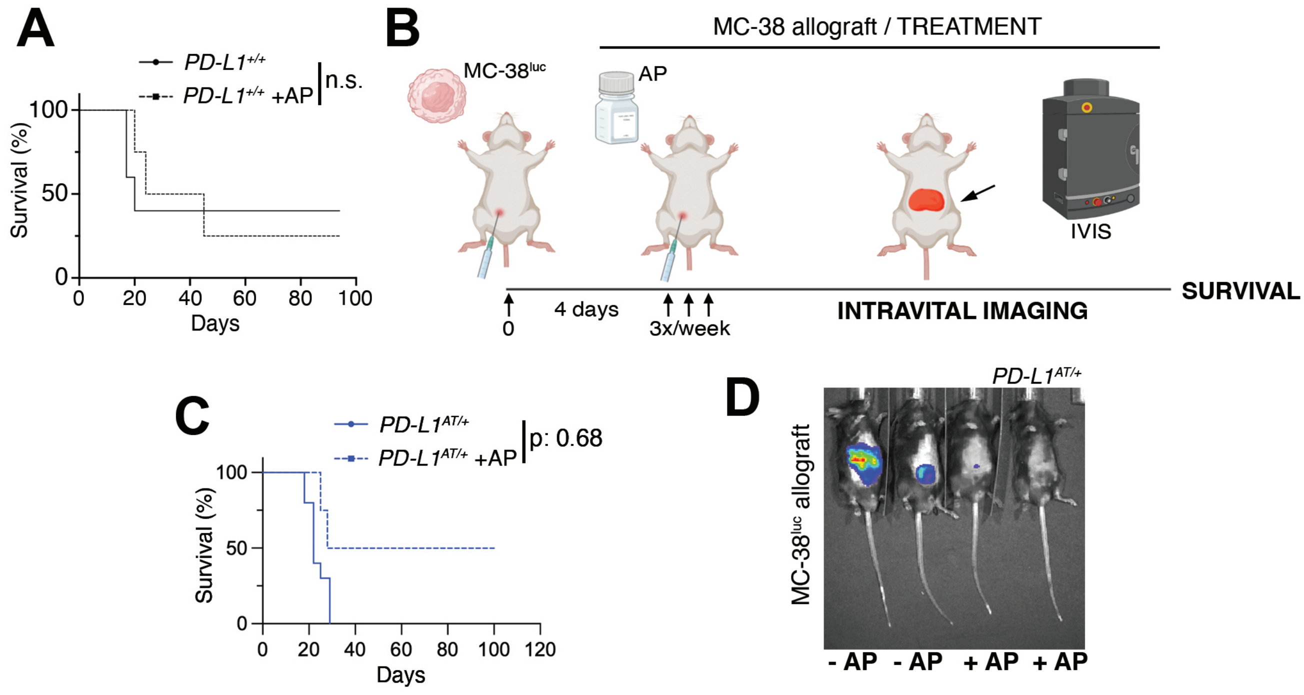

**Figure S4. Impact of depleting PD-L1$^{+}$ cells in MC-38$^{luc}$ allografts.** (A) Kaplan-Meier survival curve of control and AP20187-pretreated *PD-L1*$^{+/+}$ mice after i.p. inoculation of MC-38$^{luc}$ allografts. The p value was calculated with the Mantel-Cox log rank test. n.s.: non-significant. (B) Schematic overview of the treatment experimental workflow. $5 \cdot 10^5$ MC-38$^{luc}$ cells were intraperitoneally injected into mice. 4 days later mice were injected i.p. with AP20187 (2.5 mg/kg) 3 times a week for the duration of the experiment. (C) Kaplan-Meier survival curve of control and AP20187-pretreated *PD-L1*$^{AT/+}$ mice after i.p. inoculation of MC-38$^{luc}$ allografts in the treatment model. The p value was calculated with the Mantel-Cox log rank test. (D) Representative IVIS image of mice from the experiment defined in (C).

### EXTENDED METHODS

#### Cell culture

293T cells (ATCC) were grown in DMEM supplemented with 10% FBS and 1% Penicillin/Streptomycin and transfected in media containing no antibiotics. MC-38 cells (kind gift from Eduard Batllé) were cultured in DMEM supplemented with 10% FBS and 1% Penicillin/Streptomycin. For the generation of an MC-38 clone expressing luciferase, MC-38 cells were transduced with "pLenti CMV B5-Luc Blast" lentiviruses (Addgene, #21474). MEFs isolated from 13.5 d.p.c. embryos were cultured in DMEM (Sigma, D5796) supplemented with 10%-15% FBS (Sigma) and 1% Penicillin/Streptomycin (Gibco, #11548876) in low-oxygen conditions. Cells were passaged every 3 or 4 days in a 1:4 ratio. IFNγ (Peprotech, #315-05) was added to culture media at 10, 50 or 100 ng/ml concentration. For the experiments assessing the effect of AP20187, MEFs were plated in DMEM containing 10% FBS for 2 days in the presence or absence of IFNγ (10 ng/ml). To evaluate the effect of the drug in non-growing cells, MEF were washed twice with PBS and grown in low-serum media (0.1% FBS) for 3 days in the presence or absence of IFNγ. In all cases, AP20187 was added to culture media at a final concentration of 100 nM for the last 3 days. Resting splenic B cells were isolated and cultured in IFNγ (10 ng/ml), LPS (10 ng/ml) (Sigma, L2630) and M-CSF (10 ng/ml) for 24 hours before exposition to AP20187 (100 nM) with or without the caspase inhibitor I (20 µM) (Sigma-Aldrich, #627610) for 24 hours.

#### Lentiviral infection

Lentiviruses were produced by transfection of 293T cells with lentiviral transfer vectors and the packaging plasmids pMDL, pRev and VsVg at a 1:0.65:0.25:0.35 ratio. Transfection was performed using Lipofectamine 2000 (ThermoFisher, #11668019) and Opti-MEM Reduced Serum Medium (Gibco, #31985070) as recommended by the manufacturer. Viral supernatants were collected 48 h following transfection, filtered through a 0.45 µm filter and either added to target cells or frozen at -80ºC.

**CRISPR activation**

In order to obtain SAM-compatible mESC, PD-L1$^{AT/+}$ mESC were simultaneously infected with lentiviruses coding for MS2-p65-HSF1 and dCas9-VP64 [1]. Two days after infection, cells were selected with 10 µg/ml blasticidin (Invitrogen) and 200 µg/ml hygromycin B (Calbiochem). Selected clones were transduced with lentiviruses expressing a sgRNA targeting the *Cd274* promoter (5´-CACCGTTCGGTTTCACAGACAGCGG-3´) and EGFP expression was evaluated by flow cytometry.

**Immunohistochemistry**

Tissues were fixed in formalin and embedded in paraffin for subsequent processing. Consecutive 2.5-µm sections were treated with citrate for antigenic recovery and processed for immunohistochemistry with antibodies against EGFP (Cell signaling, #2956), mPD-L1 (Abcam, ab213480), CD8 (CNIO Monoclonal Antibody Unit, clone OTO94A) and CD45 (BD Biosciences, #553089). IHCs were scanned and digitalized with a MIRAX system (Zeiss) for further analyses.

**Flow cytometry**

Cells were trypsinized, pelleted and resuspended in PBS with DAPI. For experiments using AP20187, culture media and the PBS from the washing steps were also collected to include dead cells in the analyses. Cells were blocked with PBS +1% BSA (Roche #10735086001) for 30 minutes and stained with an anti-mPD-L1 antibody antibody (Biolegend, 124313).. mPD-L1 and EGFP expression were then evaluated by flow cytometry with a FACSCanto II (BD Biosciences), and data analysed with FlowJo (BD Biosciences).

**Immunoblotting**

For WB analyses, cells were washed once with PBS, and lysed in RIPA buffer (Tris-HCl 50 mM, pH 7.4, NP-40 1%, Na-deoxycholate 0.25%, NaCl: 150 mM, EDTA 1 mM) containing protease and phosphatase inhibitors (Sigma). Samples were resolved by SDS-PAGE and analyzed by standard WB techniques. Primary antibodies against GFP, PD-L1, GAPDH and β-ACTIN were used (see the list at the end of the Methods section with the references to the antibodies). Protein blot analyses were performed on the LICOR platform (Biosciences).

### *In vitro* immunofluorescence

MEFs were grown on a IbiTreat µSlide 8 well plate (Ibidi, #80826) and fixed with 1% PFA (EMS, #15710). Cells were blocked with PBS containing 1% BSA (Roche, #10735086001) for 30 minutes at RT and incubated with anti-mPD-L1 primary antibody (CNIO Monoclonal Antibody Unit, clone GOYA536A) in PBS for 1 hour at RT and Goat anti-Rat IgG (H+L) Alexa Fluor 594 (Invitrogen, A11007) in PBS for 1 hour at RT. Nuclei were counterstained using 10 µg/ml Hoechst 33342 (Invitrogen, H3570) in 2x SSC for 30 minutes at RT. Images were acquired using a SP8 microscope (Leica) with a 63x magnification lens at non-saturating conditions. Images were then processed with ImageJ.

### Tissue immunofluorescence

Immunofluorescence on tissue sections was carried out as previously described[2]. Briefly, after antigen retrieval, the sections are rinsed, and permeabilised in PBS containing 0.25% Triton X100 and 0.2% gelatine. After that, they were blocked in 5% BSA in permeabilisation buffer. Sections were then incubated with the primary antibodies in 1% BSA overnight at room temperature, and with the secondary antibodies for 1 h at room temperature. Finally, after rinsing, the sections are incubated in 10 mM CuSO4/50 mM NH4Cl solution, dried, and mounted with ProLong Gold antifade (ThemoFisher, P10144) mounting media. Images were captured with a Leica SP5 WLL confocal microscope. A 40x magnification lens was used and images were taken at non-saturating conditions.

### Anti-mouse PD-L1 mAb

A new anti-mouse PD-L-1 mAb (clone GOYA536A) was produced by immunizing Wistar rats with HEK293-expressed extracellular domain (ec) of PD-L1 fused to Fc fragment. Wistar rats (Charles River Laboratories, France) were injected intraperitoneally (three times at 14-day intervals) with 100µg of ecPDL1-Fc and Complete Freund's adjuvant (Difco). A 150µg last booster of the recombinant ecPD-L1-Fc protein was injected intraperitoneally and splenocytes were isolated and fused 3 days later. Hybridoma supernatants were screened by ELISA using HEK293T cells transfected with the PCMV6-mPDL1-MYC-DDK plasmid (Origene, #MR203953). The rat mAb that was raised against mouse PD-L1 (clone GOYA536A) was cloned by the limiting dilution technique. All animal experiments were performed under the experimental protocol approved by the Institutional Committee for Care and Use of Animals from Consejería de Medio Ambiente y Ordenación del Territorio of the Comunidad de Madrid (Madrid, Spain) with reference number PROEX62.3/20. All efforts were made to minimize animal suffering.

**ELISA**

IL-6 levels were quantified by an anti mIL-6 ELISA kit (Invitrogen #88-7064-22) following manufacturer's instructions. Plates were analyzed using a Victor microplate reader (Perkin Elmer).

**Droplet based single-cell mRNA sequencing**. Peritoneal cavity cells were obtained by peritoneal lavage following standard procedures. Cells were collected in cold PBS containing 5% FBS and 1 mM EDTA to preserve cell viability. Cell suspension was washed with PBS containing BSA and filtered through a 40 µm cell strainer. Viable cells were enriched by magnetic separation using the Dead Cell Removal Kit (Miltenyi Biotec #130-090-101). Isolated cells were next washed and finally suspended in PBS-0.04% ultrapure BSA (Invitrogen #AM2616) at 1000 cells/µl, and shown to have a viability higher than 95% by Trypan Blue exclusion (Gibco #15250061). Approximately $10^4$ cells from each sample were loaded onto a 10X Chromium Single Cell Controller chip B (10x Genomics) following manufacturer's instructions (Chromium Single Cell 3′ GEM, Library & Gel Bead Kit v3, PN-1000075). Generation of gel beads in emulsion (GEMs), barcoding, GEM-RT clean up, cDNA amplification and library construction were all performed according to manufacturer's recommendations. Libraries were sequenced in an asymmetrical pair-end format, with 28 bases for read 1 and 56 for read 2 in a NextSeq550 instrument (Illumina) with v2.5 reagent kits.

**scRNAseq data analysis**. Bcl files were converted to fastq format with cellranger mkfastq (10x Genomics), a wrapper for bcl2fastq (Illumina), and subsequently analysed with/by the bollito pipeline[3]. Reads were aligned to the Gencode mouse reference (GRCm38) and quantified using the STARsolo aligner[4]. The Seurat toolkit[5] was used to perform the cell-based quality control, normalisation, integration and clustering steps. We filtered out cells with less than 750 genes detected and high mitochondrial (>10%) and ribosomal (>40%) gene content. Cells with more than 4000 detected genes were considered doublets. Low-abundance genes, those expressed in less than 2 cells, were also removed from the dataset. A total of 14347 cells were recovered. Samples were normalised using the *sctransform* approach[6] and integrated. The first 20 components were selected to cluster the samples and a Uniform Manifold Approximation and Projection (UMAP) was applied for their visualisation. We used the SingleR[7] annotation scores to guide the annotation of each cluster. Differential gene expression was performed using Seurat's wilcox test. Genes expressed in less than 25% of the cells were filtered out. The estimated significance level (P value) was corrected to account for multiple hypotheses testing using Benjamini and Hochberg False DiscoveryRate (FDR) adjustment. Genes with FDR less than or equal to 0.05 were selected as differentially expressed. Significantly upregulated or downregulated genes were introduced into PANTHER to perform a Gene Ontology Biological Process enrichment analysis (http://geneontology.org)[8-10]. Finally, a Gene set enrichment analysis (GSEA) was performed using the *fgsea package*[11].

**Antibodies**

| Antibody | Use | Dilution | Reference |
|---|---|---|---|
| GFP | WB | 1:1000 | Cell Signaling, #2956 |
| PD-L1 | WB | 1:1000 | Abcam, ab213480 |
| GAPDH | WB | 1:1000 | Cell Signaling, #2118 |
| β-ACTIN | WB | 1:50000 | Sigma, A5441 |
| CD45R/B220 PE | FACS | 1:400 | BD Biosciences, 553089 |
| PD-L1 PE-Cy7 | FACS | 1:400 | Biolegend, 124313 |
| GFP | IHC | 1:100 | Cell signaling, #2956 |
| PD-L1 | IHC | 1:2000 | Abcam, ab213480 |
| CD45 | IHC | 1:200 | Cell signaling, #557390 |
| CD8a | IHC | 1:200 | Clone OTO94A* |
| PD-L1 | Tissue IF | 1:500 | Abcam, ab213480 |
| GFP | Tissue IF | 1:500 | Abcam, ab13970 |
| mPD-L1 | IF | 1:20 | Clone GOYA536A* |
